# Phantom epistasis through the lens of genealogies

**DOI:** 10.1101/2024.12.03.626630

**Authors:** Anastasia Ignatieva, Lino A. F. Ferreira

## Abstract

Phantom epistasis arises when, in the course of testing for gene-by-gene interactions, the omission of a causal variant with a purely additive effect on the phenotype causes the spurious inference of a significant interaction between two SNPs. This is more likely to arise when the two SNPs are in relatively close proximity, so while true epistasis between nearby variants could be commonplace, in practice there is no reliable way of telling apart true epistatic signals from false positives. By considering the causes of phantom epistasis from a genealogy-based perspective, we leverage the rich information contained within reconstructed genealogies (in the form of ancestral recombination graphs) to address this problem. We propose a novel method for explicitly quantifying the genealogical evidence that a given pairwise interaction is the result of phantom epistasis, which can be applied to pairs of SNPs regardless of the genetic distance between them. Our method uses only publicly-available data and so does not require access to the phenotypes and genotypes used for detecting interactions. Using simulations, we show that the method has excellent performance at even low genetic distances (around 0.5cM), and demonstrate its power to detect phantom epistasis using real data from previous studies. This opens up the exciting possibility of distinguishing spurious interactions in *cis* from those reflecting real biological effects.

## 1 Introduction

The study of the genetic basis of human traits and diseases has progressed rapidly in recent years. Fuelled by large biobanks, genome-wide association studies (GWASs) have led to the discovery of thousands of genetic variants associated with a wide array of phenotypes (Sollis et al., 2023; Tam et al., 2019), shedding light on their genetic architecture (Lappalainen et al., 2024; Watanabe et al., 2019) and enabling the prediction of disease risk based on genetic data through polygenic scores (PGSs) (Lambert et al., 2021; Torkamani et al., 2018).

The overwhelming majority of genetic associations identified so far involve variation at a single locus (Tam et al., 2019). Comparatively less attention has been given to gene-by-gene (G×G) interactions, or epistasis, between pairs or larger groups of mutations, where the simultaneous presence of mutations at two or more distinct loci has a combined average effect on the phenotype that is different from the sum of their individual effects in isolation (Cordell, 2009; Phillips, 2008). It is essential to differentiate functional epistasis, which describes the underlying mechanistic interactions between genes at a biochemical or cellular level, from the quantitative genetic notion of statistical epistasis. The latter, at a population level, measures the deviation from additivity in a statistical model of allelic effects on a phenotype. The magnitude of this statistical epistasis, and the resulting epistatic variance component, is fundamentally dependent on the frequencies of the interacting alleles within the population. Consequently, functional epistasis can be both strong and widespread, yet contribute very little to the total genetic variance if the variants (or, more precisely, the haplotypes carrying the interacting combination of alleles) are rare; this problem is exacerbated by the fact that the frequency of an interaction is dependent on the product of the frequencies of the individual variants, making it significantly lower than either frequency. Moreover, estimates of the additive effect of variants will tend to capture part of the effects of their possible interactions due to standard properties of linear regression. Low epistatic variance in a population therefore does not preclude the existence of significant functional gene interactions. In humans, few signals of statistical epistasis have been found (Lappalainen et al., 2024; Wei et al., 2014) and it has been established that this has a limited role in explaining phenotypic variance (Hill et al., 2008; Hivert et al., 2021; Sella and Barton, 2019), and so is unlikely to explain a substantial fraction of ‘missing heritability’ (Manolio et al., 2009) or allow for improved PGS performance. However, detecting a statistical interaction between two parts of the genome suggests a functional interaction between them and so could improve our understanding of the biological function underlying genetic associations (Domingo et al., 2019; Mackay, 2014; Phillips, 2008), a fundamental open problem in the field (Lappalainen et al., 2024).

The search for genetic interactions is made challenging by the vast search space of possible variant combinations which, for pairwise interactions, grows quadratically in the number of variants considered (Wei et al., 2014). This has raised obstacles along two dimensions. First, exhaustive testing for all possible pairwise combinations of variants has a large computational cost. Second, even if this computational barrier is surmounted, performing a great number of statistical tests requires a stringent multiple testing correction to avoid rampant false positives, resulting in loss of statistical power; this is exacerbated by such tests having a higher-dimensional parameter space than tests for direct effects. Multiple software tools have been developed that make exhaustive testing for epistasis possible (Upton et al., 2016), however they have not yielded much fruit, likely due to a lack of power. As an alternative, methods that first perform a strategic reduction of the search space by leveraging statistical or biological features of the loci under consideration (Sun et al., 2014) appear more promising and have enabled the identification of a few robust interactions (GAPC & WTCCC2, 2010; TASC et al., 2011; Masuda et al., 2015; Lyon et al., 2022).

Recently, a novel statistical challenge to the reliable detection of interactions has been recognised and explored. Termed ‘phantom epistasis’, this phenomenon arises when the statistical evidence for the existence of a genetic interaction fades when the *additive* effect of a third variant is considered.Phantom epistasis was first clearly identified as a problem following a study which examined possible interactions impacting transcription levels in humans: Hemani et al. (2014, retracted) considered the expression of 7,339 genes, measured through RNA sequencing, and detected 30 statistically significant interactions affecting 19 different genes that replicated in two external datasets. However, Wood et al. (2014) showed with different data that none of these interactions remained statistically significant once the most strongly associated variant with the corresponding phenotype was included in the model. A crucial difference between the datasets used in the two studies was sequencing depth: the former study used genotyping array data (∼ 530,000 single-nucleotide polymorphisms, SNPs) while the latter used a whole-genome sequencing (WGS) dataset. Wood et al.’s findings suggest that the interactions originally reported were, in effect, tagging a third variant whose effect on the phenotype was fully additive (and which was not included in the data of Hemani et al.).

More recently, de los Campos et al. (2019) have demonstrated, using a simplified statistical model of genetic interactions, that phantom epistasis can arise as the result of mutual but imperfect linkage disequilibrium (LD) between the two spuriously interacting SNPs and a third, unobserved, causal quantitative trait locus (QTL) which has a fully additive effect. They state that it is not possible to empirically test for this condition, as the QTL genotype is unknown. This phenomenon is a direct statistical consequence of third-order LD, the non-random association of alleles at three loci (Bennett, 1954). In this scenario, the interaction term becomes a proxy for the unobserved causal variant due to this higher-order dependency. The potential for this to create phantom epistasis is not merely theoretical: for instance, Zan et al. (2018) found higher-order LD to be prevalent in *Arabidopsis thaliana* sequencing data. Evolutionary forces such as selection can introduce and maintain such higher-order associations (Grote et al., 1998), potentially creating complex signals of phantom epistasis around functionally-important genomic regions. Hemani et al. (2021) reproduced spurious interactions in simulations and recommend adjusting for fine-mapped additive effects when testing for epistasis. Thus, the challenge posed by phantom epistasis remains unsolved, with the best solution currently available being to carefully account for observed additive effects, with no guarantee that this will be sufficient to avoid false-positive interactions. Using WGS data does largely resolve the problem of missing causal variants, although not entirely, since this will still miss structural variants and SNPs appearing in repetitive regions which are difficult to sequence.

In this article, we develop a method that can provide evidence against the existence of an unobserved additive effect which could explain a putative interaction. Our key observation is that, while we cannot (by definition) observe a missing genetic variant, it is possible to calculate a proxy genotype with which any such unobserved QTL would have to be highly correlated for phantom epistasis to emerge, and to search for evidence of its presence – or of its absence – within a genealogical framework. We do this by considering the conditions for phantom epistasis in terms of conditions on the existence of clades of samples within the underlying genealogy. For instance, if the set of samples carrying both of the detected interacting SNPs itself forms a clade within the genealogy, and a variant shared by these samples is present but not included in the statistical analysis, the interaction term will act as a proxy for this unobserved additive-effect variant (likely resulting in a false positive in a test for interactions).

We present a computational tool (Spectre: Searching for Phantom EpistatiC interactions using TREes) implementing two methods of searching for evidence of phantom epistasis, which can be applied to summary statistics from association studies conducted using large-scale genotype data, and genealogies (in the form of ancestral recombination graphs, ARGs) reconstructed using a WGS dataset; we use the 1000 Genomes Project (1KGP, 1000 Genomes Project Consortium, 2015). We proceed by calculating genotype proxies for potential unobserved causal (additive effect) QTLs, and then (1) searching for clades in the genealogy that can cause phantom epistasis through being in LD with these proxies, and (2) searching for evidence *against* the existence of such problematic clades. We show that, as previously noted, phantom epistasis is relatively likely to arise when the two SNPs are in close proximity. However, our method allows to explicitly test for the presence of phantom epistasis for SNPs in *cis*, without requiring an arbitrary cut-off on the genetic distance between them as has been suggested in the literature. We use simulation studies to demonstrate the effectiveness of these methods and apply them to the interactions identified by Hemani et al. (2014), which were shown to be false positives by Wood et al. (2014). Our work presents the first practical computational tool for assessing the likelihood that detected genetic interactions are due to phantom epistasis.

Code implementing the methods is available at github.com/a-ignatieva/spectre.

## 2 Methods

### 2.1 Overview of ARGs

The genealogy of a sample of sequences can be fully encoded in the form of an ARG, which can be represented as a series of local trees, describing the local genealogy at each locus (for recent overviews, see for instance Brandt et al., 2024; Lewanski et al., 2024; Wong et al., 2024; Nielsen et al., 2025). Adjacent local trees are highly correlated, as edges tend to persist across multiple local trees before being broken up by recombination. A set of samples can form a clade in the ARG if at some genomic position the set is subtended by a branch of the corresponding local tree. Each clade has a well-defined genomic span, being an interval including all the genomic positions where, in the corresponding local tree, there is a branch subtending exactly the set of samples in the clade. This is illustrated in Figure 1. ARGs reconstructed from a given sequencing dataset thus provide not only an estimate of the age of each mutation (under the infinite sites model, which we assume throughout), but also an estimate of the genomic span of the clade of its carriers. For any given ARG, it is thus possible to traverse the local trees left-to-right, and record the genomic span of each encountered clade of samples.

**Figure 1:**
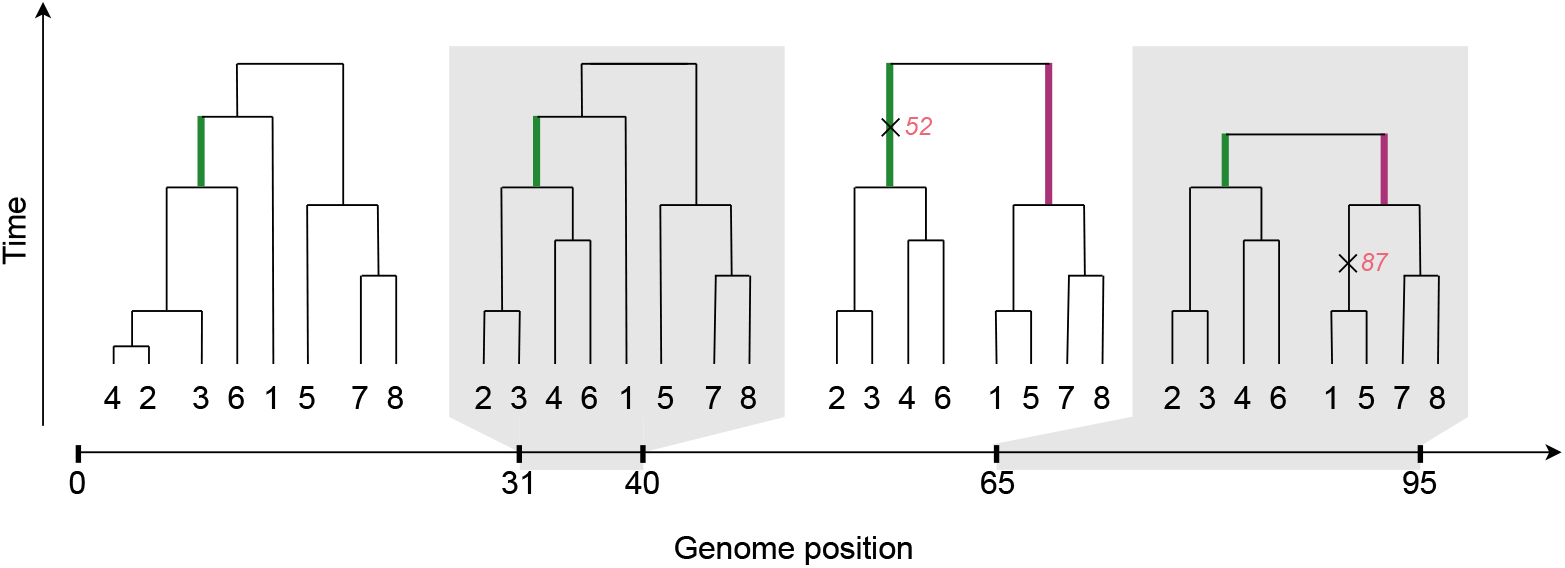
Example of an ARG with *n* = 8 samples, with four local trees shown covering the genomic region [0, 95]. The set of samples *A* ={2, 3, 4, 6} forms a clade *Â* within the entire interval [0, 95] (since there is a branch in each local tree subtending exactly this set of samples, shown in green); this clade is supported by one mutation at position 52 (shown as ×). The set of samples *B* ={1, 5, 7, 8} forms a clade 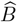 within the interval [40, 95] so has a genomic span of 55bp (in local trees 3 and 4, the branches subtending this clade are shown in purple); this clade is not supported by any mutations. The set *C* ={1, 3, 4}, for instance, does not form a clade anywhere within the shown region.

Each clade and mutation within the ARG uniquely defines a corresponding set of samples (those belonging to the clade, or carrying the mutation, respectively). A given clade 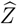 containing the set of samples *Z*, in turn, defines a corresponding (haploid) genotype vector ***z*** = (*z*_1_, …, *z*_*N*_), with *z*_*i*_ = 1 if *i* ∈ *Z* and *z*_*i*_ = 0 otherwise. Likewise, a given genotype vector ***v*** uniquely defines a corresponding set of samples *V* = {*i* : *v*_*i*_ = 1}, but it is not necessarily the case that this set forms a clade 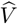 within the ARG.

### 2.2 Methods for detecting epistasis

The issue of phantom epistasis concerns only *statistical epistasis*, that is, statistical interaction terms between genetic variables in parametric linear models of phenotypic architecture (Phillips,2008). By construction, these effects are measured while averaging over all other genotypes in a population. There are two widely-used models of phenotypic architecture that include epistatic effects. Consider first the simplest linear model of a quantitative phenotype ***y*** with a single genetic interaction between two loci (with genotype vectors ***x***_1_ and ***x***_2_):

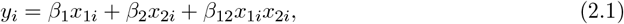

where *i* indexes the samples. The canonical test for a genetic interaction is to fit a linear regression model and perform a *t*-test of significance for the estimated *β*_12_ (Cordell, 2009) (using logistic regression when the phenotype is binary). A less parsimonious model separates additive from dominance effects and allows interactions between these two classes of effects:

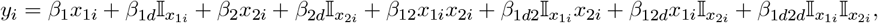

where 𝕀_*x*_ = 1 if *x* = 1 and 0 otherwise. Under this model, we can test for the existence of an epistatic effect of any of these four types (additive-by-additive, additive-by-dominance, etc.) at once through an *F* -test with four degrees of freedom.

In their original analysis of epistasis affecting expression levels, Hemani et al. (2014) performed *F* -tests with four degrees of freedom using the computational implementation of Hemani et al. (2011). In later work, they showed through simulations that such a test is prone to phantom epistasis and stated that the same is true of the *t* -test approach (Hemani et al., 2021), which is the model de los Campos et al. (2019) considered in their work. Given the high-dimensionality of the search space and the fact that very few interactions have been identified using human genetic data, we explore the issue of phantom epistasis using the simpler model (2.1). This approach makes tractable the derivation of a key result used in our method.

### 2.3 Phantom epistasis

Consider testing for an interaction between SNP1 and SNP2 (at positions *m*_1_ and *m*_2_, with genotype vectors ***x***_1_ and ***x***_2_, and sample sets *X*_1_ and *X*_2_, respectively) within the linear regression model

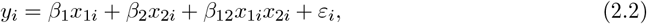

where *ε*_*i*_ is an error term. Assuming that neither the SNPs nor their interaction has a causal effect on the phenotype (such that *β*_1_ = *β*_2_ = *β*_12_ = 0), it may nevertheless be the case that our estimate of the coefficient *β*_12_ is statistically significant if a third SNP *z*_*i*_ which does have a causal effect on *y*_*i*_ is correlated with the interaction term after controlling for *x*_1*i*_ and *x*_2*i*_ (strictly, if the partial covariance of *z*_*i*_ and the interaction term conditional on *x*_1*i*_ and *x*_2*i*_ is sufficiently large relative to the partial variance of the interaction term; this follows from standard linear regression results, and a detailed derivation is given in SI, Section S1.3). We note that the same principles of LD with an unobserved causal variant giving rise to phantom epistasis can also cause phantom dominance. Specifically, a spurious dominance effect can be detected at a marker locus if it is in LD with a purely additive causal variant that has a different allele frequency.

Comparing this setting to a standard GWAS where the individual effects of SNPs on a phenotype are tested, there is both a similarity and an important difference worth noting. Whenever we test for the direct effect of a variant on a phenotype, it is understood that a statistically significant effect does not necessarily imply that the variant in question has a causal effect on the phenotype. Rather, we recognise that it may be merely correlated with such a causal effect in its vicinity, a phenomenon known as ‘tagging’, and that fine-mapping is required to attempt to pinpoint the true causal variant. In a sense, a similar claim could be made about a statistically-significant interaction. The crucial difference is that, if a third variant with an *additive* effect explains an apparent interaction signal, this changes the nature of the putative interaction effect being reported. The claim that a statistical interaction has an effect on a phenotype implies that this effect cannot be fully accounted for by any combination of simple additive effects, and that a fundamentally more complex phenomenon is at work (in other words, a purely additive model is not a fully accurate statistical representation of the genetic basis of this phenotype). If it turns out that the additive effect of a third variant can explain the interaction signal, this is no longer true.

To begin examining this issue from a genealogy-based perspective, let us assume a haploid model for simplicity and consider the following example matrix of genotypes:

| $\mathbf{x}_1$ | $\mathbf{x}_2$ | $\mathbf{s}$ | $\mathbf{z}$ |
| --- | --- | --- | --- |
| 0 | 0 | 0 | 0 |
| 1 | 0 | 0 | 0 |
| 1 | 1 | 1 | 1 |
| 1 | 0 | 0 | 0 |
| 0 | 0 | 0 | 0 |
| 1 | 1 | 1 | 1 |
| 0 | 1 | 0 | 0 |
| 0 | 1 | 0 | 0 |

where ***s*** := ***x***_1_ ∘ ***x***_2_ (‘∘’ denoting element-wise product) represents the simultaneous occurrence of the two mutations, and ***z*** is a third mutation that happens to be perfectly correlated with ***s***. If ***z*** is the true causal variant for the phenotype but only ***x***_1_, ***x***_2_ and ***s*** are included in the model, ***s*** will likely appear statistically significant as it is perfectly correlated with the true (additive) causal effect, when in fact no epistasis is present.

We note that variants like ***z*** are not the only kind that can lead to spurious interactions. Suppose that a significant interaction is detected between ***x***_1_ and ***x***_2_. Then define the four ‘target sets’ as

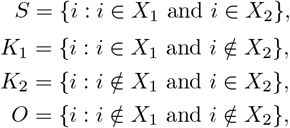

where *X*_1_ = {*i* : *x*_1*i*_ = 1} and *X*_2_ = {*i* : *x*_2*i*_ = 1}. Phantom epistasis can arise if any of these sets forms a clade in the ARG (at *any* position along the genome), but no SNP corresponding to this clade is included in the model (implying that a causal variant shared by the samples in this clade might exist but has not been sequenced). This is because inclusion of such a SNP would mean that the interaction term can be represented by a linear combination of ***x***_1_, ***x***_2_, and the genotype vector of this SNP. For instance, consider the following matrix corresponding to the ARG in Figure 2:

**Figure 2:**
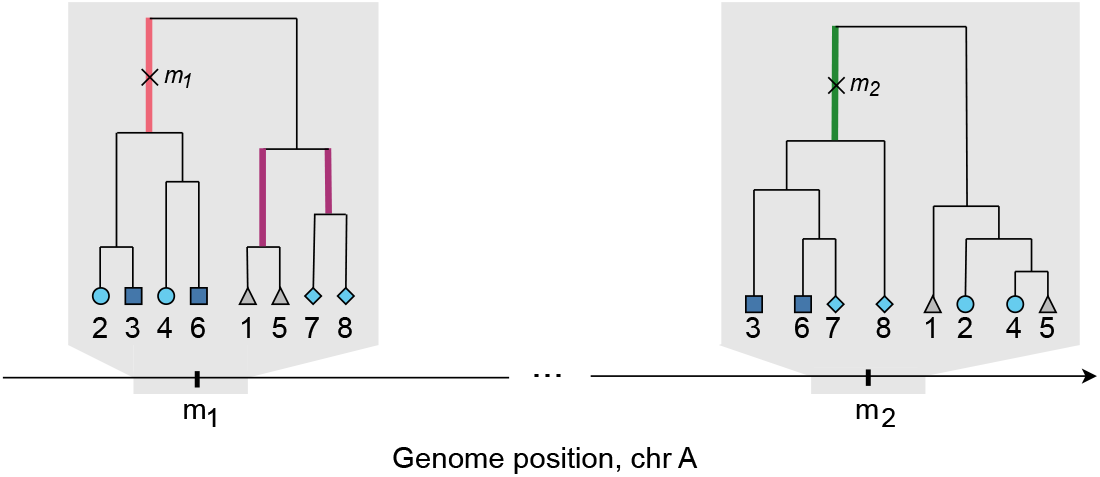
The two mutation events, SNP1 at *m*_1_ and SNP2 at *m*_2_, are shown as × on the corresponding branches (in red and green) of the local trees at *m*_1_ and *m*_2_, respectively. Here *X*_1_ = {2, 3, 4, 6}, *X*_2_ = {3, 6, 7, 8}, and the corresponding target sets are *S* = {3, 6} (dark blue squares), *K*_1_ = {2, 4} (light blue circles), *K*_2_ ={7, 8} (light blue diamonds), *O* = {1, 5} (grey triangles). In these two trees the set *S* does not form a clade, since there is no branch that subtends only the samples in *S*; the same is true for set *K*_1_. The sets *K*_2_ and *O* form clades in the local tree at *m*_1_; branches subtending these clades are shown in purple. Thus, if the causal mutation occurs on one of the purple branches but is not sequenced, phantom epistasis could arise and result in a significant interaction term between ***x***_1_ and ***x***_2_.

| $\mathbf{x}_1$ | $\mathbf{x}_2$ | $\mathbf{s}$ | $\mathbf{k}_1$ | $\mathbf{k}_2$ | $\mathbf{o}$ |
| --- | --- | --- | --- | --- | --- |
| 0 | 0 | 0 | 0 | 0 | 1 |
| 1 | 0 | 0 | 1 | 0 | 0 |
| 1 | 1 | 1 | 0 | 0 | 0 |
| 1 | 0 | 0 | 1 | 0 | 0 |
| 0 | 0 | 0 | 0 | 0 | 1 |
| 1 | 1 | 1 | 0 | 0 | 0 |
| 0 | 1 | 0 | 0 | 1 | 0 |
| 0 | 1 | 0 | 0 | 1 | 0 |

The interaction term can be written equivalently as

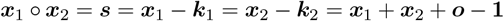

(where **1** is a vector of 1’s), so inclusion of any of ***s, k***_1_, ***k***_2_, ***o*** would result in the model being unidentifiable. On the other hand, if the genotype of the true causal variant is highly correlated with ***s, k***_1_, ***k***_2_ or ***o*** conditional on ***x***_1_, ***x***_2_, omitting this variant from the model will cause spurious inference of a significant interaction effect.

Phantom epistasis can thus arise with the existence of any clade 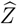 such that the corresponding set of samples *Z* is highly enough correlated (positively or negatively) with a target set, after conditioning on ***x***_1_ and ***x***_2_. It is important to note that while these four target sets are linearly dependent once the main effects are accounted for (meaning the statistical test for the interaction itself is one-dimensional), we consider each one as a distinct genealogical scenario. A phantom signal could arise from an unobserved causal variant being correlated with the specific haplotype partition corresponding to any of these sets, so we check all four possibilities.

Note that, theoretically, it does not matter where along the genome 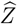 arises. Under the neutral panmictic coalescent model, the probability that a set of |*Z*| samples forms a clade in a random local tree of size *n* can be easily calculated (SI, Section S1.2); this is negligible unless |*Z*| is small, or the required level of correlation with a target set is low. The probability of 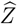 existing is, however, generally elevated at genomic positions near *m*_1_ and near *m*_2_: for instance, since *S* = *X*_1_ ∩ *X*_2_ is a subset of *X*_1_, near *m*_1_ we need to instead consider the probability that *Z* forms a sub-clade of a clade of size |*X*_1_|, which is higher than the probability that *Z* forms a clade in a tree of size *n*. Note that this is true even when *m*_1_ and *m*_2_ are very far apart. When *m*_1_ and *m*_2_ are close together, however, we know that *S* is a subset of *X*_1_ and a subset of *X*_2_, which both form clades in the region [*m*_1_, *m*_2_], so the probability of interest will be further elevated. This reasoning aligns with phantom epistasis being more likely to affect SNPs nearby in *cis*; it further implies that, for each target set, we only need to consider the probability of a highly-correlated clade existing at genomic positions near *m*_1_ and *m*_2_.

We note that population structure, through its effects on shared ancestry across the genome, makes it more likely that a specific set of samples (corresponding to a subpopulation) will form a clade at any given locus. The resulting non-gametic phase disequilibrium at large distances can be of the same order of magnitude as gametic phase disequilibrium, and both can theoretically cause phantom epistasis. Both of these sources of long-range correlation are accounted for when estimating effect sizes, however, by incorporating principal components of the genetic relationship matrix as covariates in the model or by employing linear mixed models. These corrections for population structure control for the underlying ancestral confounding that generates them. Our method focuses instead on analysing the remaining localised signals between variants nearby on the same chromosome.

### 2.4 Genealogy-based testing for phantom epistasis

Typically, interaction tests are performed using large (unphased) genetic datasets, and hence it is difficult to calculate the target sets for each pair of tested SNPs as above, and not possible to directly estimate the probability that they form clades. We thus take the following approach. For each pair of SNPs with a significant interaction term in an association study, we calculate the corresponding target sets using an ARG reconstructed from WGS data from the same population as (but not necessarily containing samples from) the study cohort from interaction testing. We define each clade by its set of descendant samples, and traverse the ARG, recording the genomic position at which each observed clade first arose, and that at which it disappeared (due to recombination).

We propose two complementary tests to assess the chance of phantom epistasis, each providing a different type of evidence. The two tests are designed to work in tandem: one actively searches for positive evidence of a phantom effect, while the other quantifies the evidence against one. Our first test (Test 1) is a direct search for a potential causal variant. This searches for specific clades in the ARG that are sufficiently correlated with the interaction’s target sets to be a plausible driver of the observed signal. This test is most powerful when it identifies a specific, high-confidence candidate clade (or clades) that could explain away the interaction. However, a failure to find a causal clade with Test 1 is not definitive proof that the interaction is real. Therefore, our second test (Test 2) approaches the problem from the opposite direction by quantifying the residual uncertainty. It evaluates the local genealogy to identify regions where the existence of a phantom epistasis–causing variant cannot be confidently ruled out. The two tests are therefore complementary. The strongest evidence for phantom epistasis arises when Test 1 identifies specific candidate clades that reside within the regions found by Test 2. Conversely, the strongest evidence against phantom epistasis (and thus in favour of a true biological interaction) is when Test 1 finds no plausible candidate clades and Test 2 does not find regions where phantom epistasis is likely to arise.

#### 2.4.1 Test 1: Searching for additive effects not included in the model

Our first method of checking for phantom epistasis is by using the 1KGP ARG to check for clades (within ±1cM of each SNP) which, if excluded from the regression model, could cause a significant (but phantom) interaction to arise or, from a different perspective, could cause a significant interaction term to cease to be significant if included.

As detailed in SI, Sections S1.3 and S1.4, we build on the derivations of de los Campos et al. (2019) and on standard results for linear regression to derive the asymptotic distribution of the test statistic of the estimated coefficient 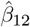 in the regression model (2.2) in a statistical test of the null hypothesis *β*_12_ = 0 under the assumption that there is no interaction but there is a correlated third variant *z*_*i*_ with a non-zero effect *b* on the phenotype (*b* is in units of standard deviations of the phenotype). Based on this we can compute, for a given significance level *α* of the original test for interaction (which yields a threshold *C* := Φ^−1^(1 − *α/*2)) and an assumed effect *b* of the variant *z*_*i*_, the probability that the test statistic exceeds the threshold *C* in absolute value, such that the null hypothesis is (incorrectly) rejected in a two-sided test. Defining an auxiliary variable *s*_*i*_ := *x*_1*i*_*x*_2*i*_ with corresponding coefficient *β*_*s*_, this is given by

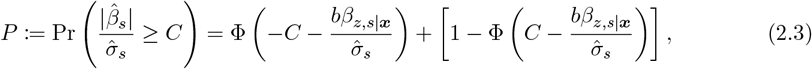

where 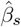 is the estimate of *β*_*s*_, 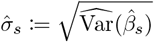 its standard error and *β*_*z,s*|***x***_ the true coefficient of *s*_*i*_ in a population regression of *z*_*i*_ on *x*_1*i*_, *x*_2*i*_ and *s*_*i*_ (Figure 3). This last coefficient depends on the partial covariance of *s*_*i*_ and *z*_*i*_ conditional on *x*_1*i*_, *x*_2*i*_ and can be readily estimated from the data used to build the ARG. And as we show in SI, Section S1.3.6, 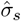 can be accurately approximated for polygenic phenotypes using only the genotype data in the same dataset.

**Figure 3:**
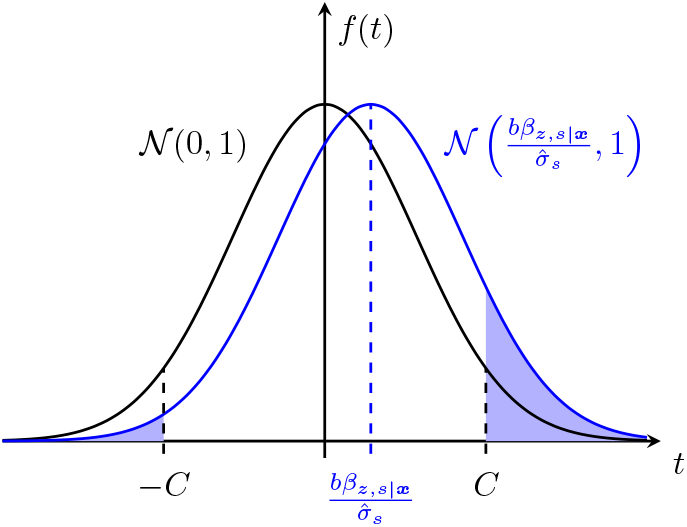
Black curve: standard Normal density of test statistic under the null; blue curve: Normal density of test statistic in the presence of an omitted variant *z*_*i*_, with mean 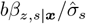 and unit variance. Shaded area shows probability of incorrectly rejecting the null hypothesis *β*_*s*_ = 0 in a two-sided test with significance level *α*, where *C* := Φ^*−*1^(1−*α/*2). Test 1A calculates *b* which makes the shaded area equal to a set probability *P*.

However, *b* is fundamentally unknowable in practice as *z*_*i*_ is unobserved. So we reframe (2.3) by instead setting a range of probabilities *P* of rejecting the null, and for each clade in the ARG we calculate the corresponding values of *b* that yield exactly those probabilities (with larger absolute values of *b* leading to greater probabilities of rejection). These values are reported by our software to aid the user in assessing how likely a given interaction is to be a false positive: if there is a third variant which need only have a small effect size to produce a large probability of incorrectly rejecting the null, this warrants caution in the confidence attached to that interaction. We refer to this procedure as Test 1A, and the idea is illustrated in Figure 3.

We also implement an alternative approach based on (2.3). Suppose that there is a third variant *z*_*i*_ with effect size *b* leading to bias in the estimated coefficient 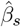 such that this coefficient converges to *bβ*_*z,s*|***x***_ (SI, Section S1.3.5). We now consider testing the non-zero null hypothesis that *β*_*s*_ = *bβ*_*z,s*|***x***_, i.e., testing whether the estimated effect of *s*_*i*_ is larger than would be expected if this interaction had no real effect on the phenotype and were only tagging the effect of the unobserved third variant *z*_*i*_. For instance, given a positive observed value for the test statistic, and an estimated *β*_*z,s*|***x***_ for a certain clade, we compute the value of *b* that would make the test statistic fall exactly at the boundary of the upper rejection region, so that the adjusted null would be rejected (just) in a two-sided test with the same significance level as the original test (Figure 4). This gives, for each clade, a threshold for *b* above which the interaction would cease to be significant. If this value for *b*, across all clades, is large, we will be more confident in that interaction as it can ‘survive’ the presence of an unobserved variant with an effect up to that size. We refer to this procedure as Test 1B.

**Figure 4:**
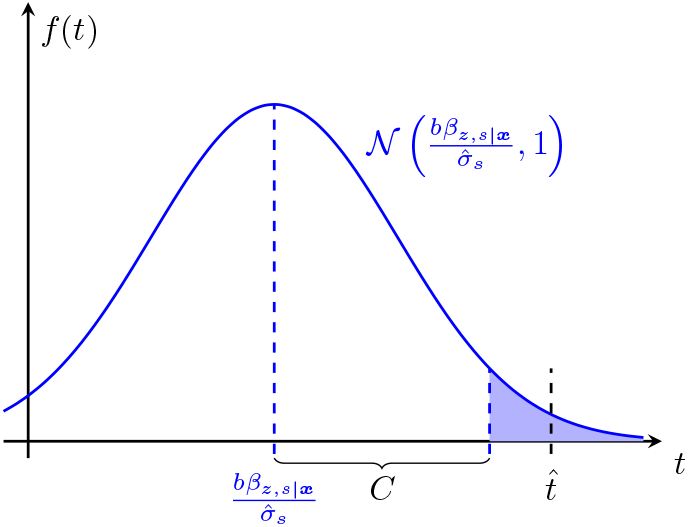
Blue curve: density of the test statistic in the presence of an omitted variant; 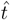 is the value of the test statistic in the original test for interactions; C is the (two-sided) critical value. Test 1B calculates *b* such that 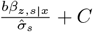 aligns with 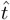.

#### 2.4.2 Test 2: Quantifying evidence against existence of unobserved additive effects

While the approach described above will identify observed sequenced variants (and those potentially shared by samples in inferred clades) which can be the cause of spuriously-inferred significant interactions, it cannot not completely rule out the possibility that such a variant exists. This can be the case if, for instance, the causal variant is a structural variant, or is present in a repeat-heavy region which is difficult to sequence, or simply due to the unavoidable presence of topological uncertainty in ARG reconstruction (Hayman et al., 2023).

We thus propose a method to consider each clade in the ARG and, within its genomic span, quantify the evidence it provides *against* the possibility of an omitted variant that causes the interaction term to have a significant effect size. To illustrate the basic idea, notice that in Figure 2 the fact that the set {2, 3} forms a clade within the genomic span of the local tree at *m*_1_ provides evidence against the possibility of the set {3, 6} forming a clade within this span. Using similar reasoning, for each target set *G*, we consider in turn each clade *Â* of the ARG (within ±1cM of each SNP), and check whether the existence of *Â* disproves the existence of all other possible clades 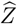 that are highly correlated with *G* and provide a sufficiently high probability of falsely rejecting the null hypothesis (or, conversely, give a sufficiently small *b* yielding such a probability) (SI, Section S1.5). Thus, at each position along the genome, we estimate the minimum value of the effect size that a hidden variant would need to have to result in a significant (phantom) interaction. This is a heuristic procedure since clearly not all possible clades are considered (since there would be an astronomical number of these, for even small ARGs), rather we make a simplifying assumption and only consider possible clades with a high direct correlation with *G*. The point of this procedure (which we call Test 2) is to narrow down the particular regions of the genome where phantom epistasis is more likely.

In summary, the method calculates the minimum (absolute) value of the effect size that a hidden variant would need to have for phantom epistasis to explain the observed data, which can then be compared against a reasonable upper bound for the absolute value of the effect size for the given trait of interest. This is done by using the clades that already exist in the ARG (Tests 1A and 1B), as well as clades that could feasibly exist but might not be present in the single reconstructed ARG being analysed due to reconstruction uncertainty (Test 2).

## 3 Results

### 3.1 Simulation studies

We simulated ARGs for 20,503 European individuals using stdpopsim (Adrion et al., 2020) (using the ‘AmericanAdmixture 4B18’ demographic model, the chromosome 22 HapMapII GRCh37 recombination map, and the default mutation rate parameter of 2.36 · 10^−8^ per bp per generation). For 10,000 of the individuals, we first simulated 100 cases of phantom epistasis. In each case we simulated a quantitative phenotype with one causal variant (true effect size 0.2 and heritability *h*^2^ = 0.01), masked this variant, and tested for interactions in the surrounding 400kb region. We set the significance threshold at 0.05, applying a Bonferroni correction based on the total number of SNP pairs after LD pruning (using Plink (Purcell et al., 2007) with window size 50, step size 5, *r*^2^ threshold 0.7). For the significant hits (which we successfully replicated using a disjoint sample of 10,000 other individuals from the same simulation) we then ran Spectre and recorded the minimum value of *b* for Tests 1A and 1B (for Test 1A, we set *P* = 0.5). We did this using the simulated ARG for the held-out 503 individuals (the same number of individuals as in the 1KGP ARG subset to European samples), as well as the ARG reconstructed for that data using Relate. We repeated this for causal SNPs of varying frequency (from 0.01 to 0.25). Next, with the same data, we simulated true interactions by selecting SNP pairs a distance *R* apart (with *R* ∈ {100kb, 500kb, 5Mb}), such that the frequency of haplotypes carrying both SNPs varied between 0.01 and 0.25. We then simulated a purely-epistatic phenotype (interaction effect size 0.2 and heritability *h*^2^ = 0.01, 0 main effects for SNP1 and SNP2), carried out regression testing (using a Bonferroni-corrected threshold) and ran Spectre as above. The code used to run the simulations and analyse the output is available at github.com/a-ignatieva/spectre-paper.

Figure 5 shows the resulting ROC curves (obtained by varying the value of the upper bound for a ‘reasonable’ effect size). Both tests demonstrate excellent performance for variants with allele frequency greater than 0.01, and the method can accurately discriminate between true and phantom epistasis for variants only 500kb apart. We observe a very small reduction in performance for reconstructed versus true ARGs, since the values of *b* calculated using true and Relate ARGs are very close (SI, Figure S3). This is as can be expected, since we only condition on the topology of the ARGs (and not, explicitly, on the event times, which are much more difficult to estimate).

**Figure 5:**
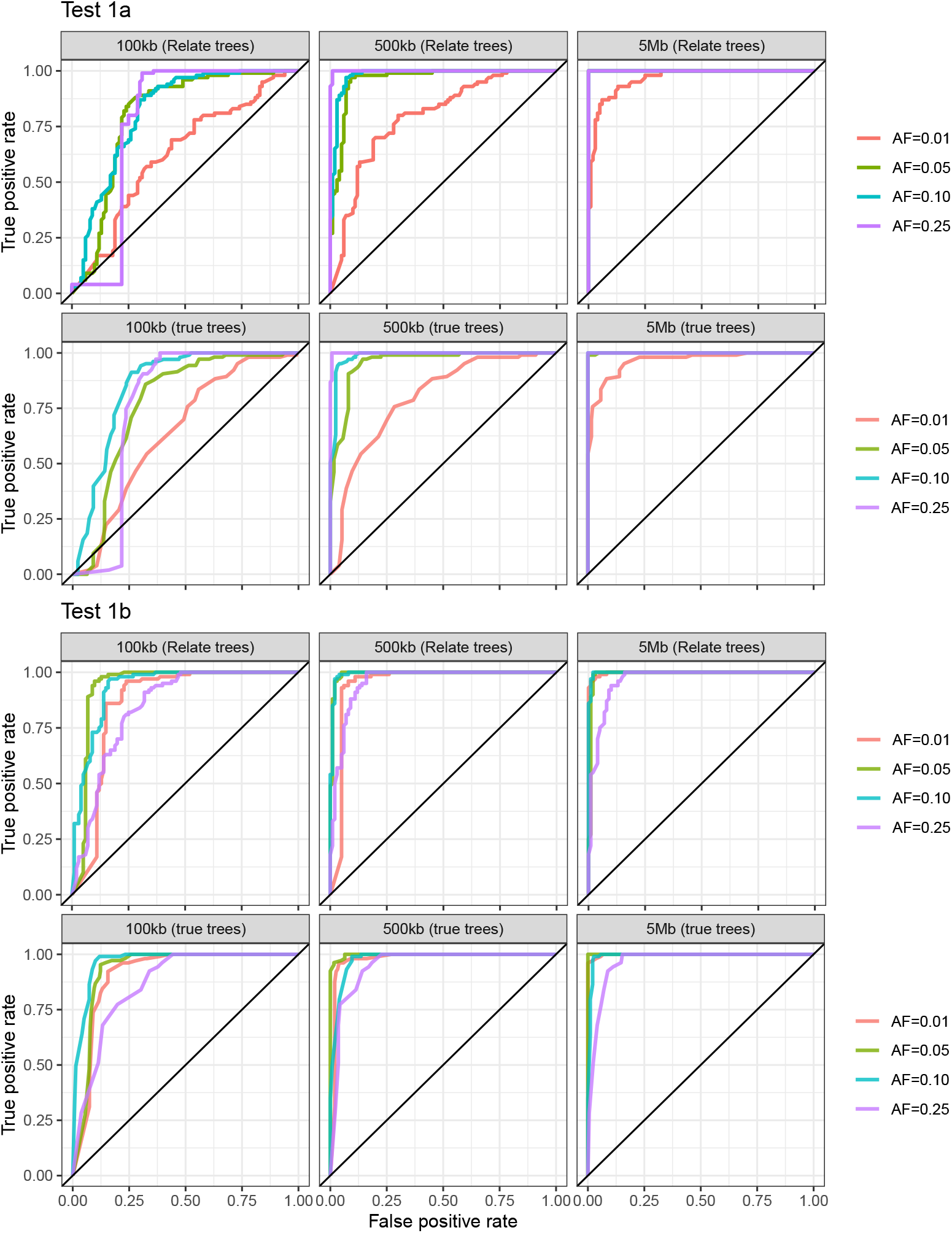
Simulation results using simulated data, with 10,000 individuals for interaction testing, and 503 individuals in the ARG. Top two rows: ROC curve for Spectre results for Test 1A, using ARG reconstructed using Relate (top row) and the simulated ARG (second row). Bottom two rows: ROC curve for Spectre results for Test 1B. For simulations of phantom epistasis, colours show allele frequency of the true causal omitted variant. For simulations of true epistasis, colours show frequency of the overlap between the two SNPs, and columns correspond to different distances between the two interacting SNPs.

We note that when our method detects phantom epistasis, it is not possible to state in these cases that the significant interaction is a ‘false positive’ as such, but rather just that it is not possible to distinguish whether epistatic or additive effects are present (due to the fundamental unidentifiability of the model when a variant exists whose set of carriers is highly correlated with a target set). As expected, this decays relatively quickly with increasing genetic distance, and (unless the two SNPs have very few carriers, relative to the size of the ARG) the probability of significant clades arising by chance is relatively low even for SNPs that are 0.5cM apart. If one or both SNPs have relatively low frequency, however, phantom epistasis can be detected even at large distances, through the existence of significant clades by random chance near one of the SNPs. Note also that the allele frequency at which phantom epistasis can be detected depends on the size of the ARG: larger ARGs mean fewer false inferences of phantom epistasis for rare variants.

### 3.2 Interactions from Hemani et al. (2014)

In order to examine the power of our method applied to real data, we consider the interactions reported by Hemani et al. (2014), which were detected using data from 846 individuals. These were,through additional sequencing of 450 individuals, shown to be the result of phantom epistasis by Wood et al. (2014). We consider the 12 (out of 17) *cis* interactions that were replicated by Wood et al. at a nominal 0.05 significance threshold (since our method is not designed to discern other causes of false positives, which can be assumed to have resulted in the lack of replication). We used the ARG reconstructed using 1KGP data (Phase 3, GRCh37) with Relate (Speidel et al., 2019). We converted the ARG into tskit format (Kelleher et al., 2016; Wong et al., 2024), and subsetted this to 503 individuals from European populations to match the data used by Hemani et al. (Powell et al., 2012). We only considered SNPs that map uniquely to a branch in the corresponding local tree, and used the HapMapII GRCh37 recombination map to calculate genetic distances. For Spectre,we used 0.05 as the significance threshold and the *p*-values from Wood et al. (2014, Table 1).

**Table 1:** Results for tested interacting SNPs from Hemani et al. (2014). Columns 1 and 2: GRCh37 positions, with frequencies given in brackets. Column 3: distance between the SNPs. Columns 4–6: minimum effect sizes output by Tests 1A, 1B, and 2, respectively. Column 7: SNP giving the lowest value of *b* per Test 1A.

| SNP1<br>(Hemani et al.) | SNP2<br>(Hemani et al.) | Distance<br>(cM) | Test 1A<br>min $b$ | Test 1B<br>min $b$ | Test 2<br>min $b$ | Top SNP<br>(this paper) |
| --- | --- | --- | --- | --- | --- | --- |
| 1:203,877,662 (0.4) | 1:203,780,591 (0.7) | 0.0 | 0.08 | 0.07 | 0.15 | 1:203,805,538 |
| 3:152,234,166 (0.3) | 3:152,116,652 (0.5) | 0.0 | 0.10 | 0.14 | 0.15 | 3:152,249,794 |
| 11:88,117,962 (0.2) | 11:88,077,479 (0.5) | 0.0 | 0.08 | 0.11 | 0.15 | 11:88,070,563 |
| 18:74,747,424 (0.5) | 18:74,732,087 (0.5) | 0.0 | 0.11 | 0.00 | 0.12 | 18:74,732,472 |
| 21:45,230,974 (0.8) | 21:45,198,355 (0.7) | 0.0 | 0.07 | 0.11 | 0.16 | 21:45,201,832 |
| 8:144,663,661 (0.6) | 8:144,613,680 (0.4) | 0.1 | 0.09 | 0.12 | 0.14 | 8:144,608,018 |
| 10:76,446,305 (0.7) | 10:75,929,517 (0.6) | 0.1 | 0.11 | 0.08 | 0.16 | 10:76,248,814 |
| 17:80,890,638 (0.6) | 17:80,827,903 (0.6) | 0.1 | 0.07 | 0.18 | 0.12 | 17:80,893,992 |
| 19:19,810,050 (0.3) | 19:19,738,554 (0.5) | 0.1 | 0.14 | 0.05 | 0.16 | 19:19,672,317 |
| 19:36,268,923 (0.4) | 19:36,147,315 (0.8) | 0.1 | 0.11 | 0.15 | 0.16 | 19:36,280,047 |
| 21:48,027,084 (0.6) | 21:47,764,477 (0.2) | 0.5 | 0.10 | 0.04 | 0.16 | 21:47,760,008 |
| 21:48,063,862 (0.2) | 21:47,776,382 (0.2) | 0.6 | 0.13 | 0.11 | 0.18 | 21:48,068,809 |

An example of the results for two of the interactions is shown in Figure 6. The *y*-axis shows the absolute value of the effect size (in units of standard deviation of the phenotype) that an omitted variant would need to have for phantom epistasis to arise, for each of the Spectre tests. At increasing distance away from the two SNPs, this value is relatively high, which suggests that a significant clade is unlikely to exist. In both cases, however, there are regions around the two SNPs (highlighted by Test 2), where the required effect size drops to low values, suggesting that we cannot rule out phantom epistasis. Indeed, in both cases there is a large number of clades (many supported by SNPs) which are highly correlated with one or more of the target sets. A summary of the full results is presented in Table 1. In all cases where the SNPs are nearby in *cis*, the effect sizes are relatively low for each test, suggesting phantom epistasis cannot be ruled out.

**Figure 6:**
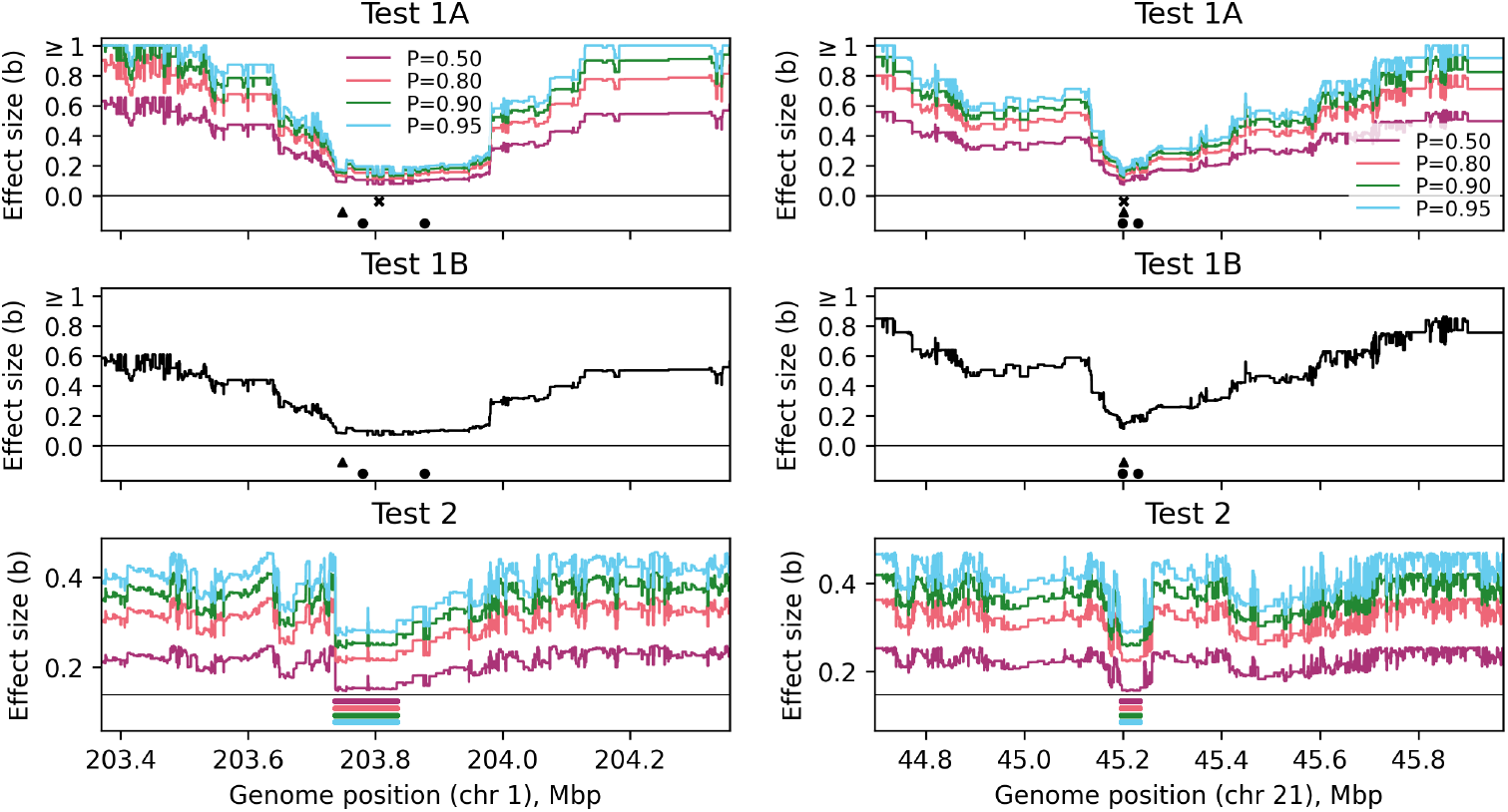
Two examples of results for interactions between SNPs in *cis* reported by Hemani et al. (2014). The *y*-axis shows the minimum effect size (in units of standard deviations of the phenotype) that an omitted variant would need to have to cause phantom epistasis, based on Tests 1A, 1B and 2. Colours show different values of the probability *P*, per (2.3); coloured horizontal bars for Test 2 show the corresponding regions where the value of *b* lies within 5% of the minimum, highlighting regions where phantom epistasis would be most likely to arise. Black points show positions of SNP1 and SNP2; black cross shows position of clade with the lowest *b* identified by Test 1A and supported by at least one SNP. Position of SNP3 from Wood et al. (2014) is shown as a triangle.

## 4 Discussion

The solutions for addressing phantom epistasis suggested in the literature so far typically involve including fine-mapped variants as covariates in the model, and ignoring pairs of variants that are in close proximity. Our method is the first quantitative approach (to our knowledge) for detecting phantom epistasis, by formulating the issue from the point of view of genealogies, and leveraging the extra information contained in reconstructed ARGs: specifically, inferred clades and their estimated genomic spans. The results can be directly interpreted as a lower bound on the effect size that an omitted variant would need to have in order to cause phantom epistasis (we note that while Test 1B requires summary statistics from the test for interactions, Tests 1A and 2 do not, and can thus be used to screen for pairs of SNPs that can be ‘safely’ tested for interaction for multiple phenotypes).

By deriving the conditions that a clade must satisfy to cause phantom epistasis, de los Campos et al. (2019) showed that it is not necessarily enough to ensure that LD between the two tested SNPs is below some threshold, or that they are far apart in genetic distance, and this is again supported by our results and simulations. Our analysis of previously-published interactions from Hemani et al. (2014) relies on the ARG inferred from the 1KGP data rather than an ARG constructed from samples in the original study. This approach is adopted out of practical necessity, with WGS data required for accurate ARG inference being typically unavailable for most study cohorts. The public 1KGP ARG serves as a high-resolution proxy for the fine-scale genealogical structure and LD patterns of the broader population from which the study samples were drawn. The validity of this cross-dataset approach hinges on the key assumption that the study cohort and the chosen 1KGP samples are well-matched and share demographic history. Under this assumption, the haplotypes (and the clades they form in the ARG) should be highly representative of those in the study cohort. However, this approach has limitations. Differences in population structure or significant genetic drift in the study cohort that is not captured by the 1KGP data could lead to differences in the underlying genealogies, potentially reducing accuracy. Further, differences in allele frequencies between the datasets could alter the composition (and size) of the target sets used for testing. These factors represent potential sources of noise that could reduce the power of our method compared to an analysis using the true, albeit unavailable, ARG which includes the study cohort. Another limitation of our approach, which is constrained by the size of the currently available ARGs, is that the tested SNPs must have relatively high frequency (to allow them to be matched to the SNPs present in the 1KGP ARG). If, for instance, there is only one sequence in the ARG which carries both interacting SNPs, by definition this sequence always forms a clade that is present in the ARG, and our tests will always detect phantom epistasis. Thus, larger ARGs need to be used to test for interactions between rarer variants. This will likely be resolved when ARGs reconstructed for biobank-scale data become available, an area of very active research (e.g. Zhang et al., 2023).

Our framework is built upon ARGs inferred under the infinite-sites model of mutation. This assumption, that every mutation occurs at a new site, is what allows each SNP to be mapped to a unique branch in the genealogy, thereby defining a distinct clade of carriers. This one-to-one mapping between a SNP and a clade is central to our approach. However, the infinite-sites assumption can be violated in reality, particularly at hypermutable sites where recurrent mutations can lead to the same variant appearing independently on different ancestral backgrounds. If the unobserved causal variant responsible for phantom epistasis arose from such recurrent mutations, it would not define a single clade in the ARG; instead, its carriers would be scattered across the genealogy, obscuring the signal our method is designed to detect and potentially reducing its power. This limitation is related to the problem of genotype imputation. The variants most likely to cause phantom epistasis are often those that are absent from genotyping arrays and poorly imputed, which can sometimes be the same variants that violate the infinite-sites model, making both their statistical imputation and their genealogical history difficult to resolve. The efficacy of our method is therefore tied to the fidelity of the underlying genealogical model. Future advances in ARG inference that can accommodate more complex mutational models will enhance the power and accuracy of this approach.

Another limitation is that our analysis is conditioned on a single point estimate of the ARG. This approach does not account for the uncertainty in topologies and clade spans inherent in genealogy reconstruction (Hayman et al., 2023; Ignatieva et al., 2025). A more robust framework would integrate our phantom epistasis tests over a posterior distribution of genealogies, for instance by using samples from a method like ARGweaver (Rasmussen et al., 2014) or SINGER (Deng et al., 2024), to ensure that the conclusions are not driven by artefacts of a single inferred genealogy. We did not pursue that approach here due to methodological and computational constraints. Our method requires ARGs inferred from large sample sizes to capture lower-frequency variants, a scale at which Bayesian ARG reconstruction tools cannot be feasibly applied. Moreover, we found in simulations that our method is very robust to ARG reconstruction error, so the impact of using a single ARG estimate is limited. We also note that within unmappable regions where the ARG cannot be reconstructed our tests cannot provide useful information. However, with the increasing availability of complete long-read sequencing datasets, and future development of genealogy reconstruction methods using such data, our tests could be used to test for phantom epistasis even in such inaccessible regions.

Our approach does not require access to the genetic and phenotypic data used in association testing, only summary statistics. It is important to stress that the method takes as input pairs of SNPs that have been detected to have significant interaction effects. It can thus only specifically detect whether these results are likely to be due to phantom epistasis as defined here (the presence of an omitted additive-effect variant). The method will not differentiate, for instance, false positives due to general inflation of test statistics or other technical issues with the test for interactions. Thus, the method does not remove the necessity of constructing sensitive and powerful tests for detecting epistasis, checking for the inflation of test statistics and replicating the results across multiple datasets. It is also important to distinguish our local fixed-effects modeling approach from the whole-genome random-effects models (e.g., linear mixed models) often used for heritability estimation or controlling for population structure. Our method is designed as a post-hoc tool to analyse a specific signal, and it intentionally uses a more localised and simpler model to directly assess the hypothesis that the signal is caused by a single nearby unobserved variant. This avoids the complexities of conditioning on the entire genome, which would alter the local covariance structure we investigate. Still regarding the differences between local statistical models and whole-genome approaches, we note that the limited impact of phantom epistasis on the estimation of the phenotypic variance explained by *cis* interactions recently reported in the literature (Smith et al., 2024) was derived under a whole-genome random-effects model. Therefore, the consequences of this phenomenon may be more severe when estimating the fixed effects of only a few variants as is the case in standard epistasis testing.

Our framework operates on phased haplotypes, and we have focused on how gametic phase disequilibrium (LD on a single haplotype) can lead to phantom epistasis. A related issue in diploid organisms is non-gametic phase disequilibrium (the correlation between alleles on homologous chromosomes) which can be of the same magnitude as gametic phase disequilibrium at large genetic distances. This phenomenon is primarily a consequence of population structure, which can create spurious associations across the genome. Standard practice in genome-wide interaction studies is to mitigate this by including principal components of the genetic relationship matrix as covariates in the model or using linear mixed models. These corrections for population structure are agnostic to whether spurious associations arise from gametic or non-gametic disequilibrium, as they control for the underlying ancestral confounding that generates them. Our method is therefore designed as a downstream tool to be applied to signals that have already been adjusted for such broad-scale confounding, allowing us to focus specifically on analysing whether the remaining signal can be explained by the fine-scale local haplotype structure captured by the ARG.

While our analysis has focused on how LD with a single, additive causal variant can generate spurious signals of epistasis between two SNPs, it is important to also consider the reverse scenario. When attempting to identify an epistatic interaction between two causal variants through SNPs that tag them, LD can have the opposite effect of masking the epistatic signal: as demonstrated by Hemani et al. (2013), the statistical power to detect true epistatic effects using markers tends to be more severely compromised by imperfect LD than the power to detect additive effects. This is because the measured epistatic interaction is dependent on the LD between both markers and their respective causal loci, leading to a more rapid decay of the detectable signal as LD weakens. This creates a significant downward bias, increasing the probability that genuine epistatic effects are underestimated or undetected. Together, these two opposing phenomena highlight that LD can not only create false positive signals of interaction but also obscure true signals, complicating efforts to resolve the genetic architecture of complex traits.

A key strength of our method is that, as noted previously in the literature, phantom epistasis appears likely to arise if the two tested SNPs are in close proximity, and significant interactions between such SNPs are typically assumed to be false positives. In general, true epistasis between nearby SNPs may well be commonplace, and the problem is only that detecting this is very difficult, in part due to the fundamental unidentifiability of the model if a variant exists whose set of carriers is correlated with a target set. Our method explicitly quantifies the genealogical evidence of this, regardless of the distance between the two SNPs, and thus offers a way of distinguishing which interactions between nearby SNPs might represent true biological effects.

An important consideration is the interpretation of clades within the inferred ARG that are not supported by any observed mutations. Intuitively, if a clade lacks supporting variants in a whole-genome sequencing panel like the 1KGP, it might seem unlikely to harbour an unobserved causal variant. However, a clade unsupported by SNPs could still tag a causal structural variant or a variant located in a genomic region that is difficult to sequence or map with short reads. These types of variants are often poorly captured in standard variant call sets but would still generate the genealogical signature our method is designed to assess. Therefore, the presence of such an unsupported but otherwise problematic clade (which our method is tailored to detect) still represents a potential source of phantom epistasis that cannot be dismissed by searching for observed SNPs in a reference panel. This benefit from using the ARG is another advantage of the method. An additional strength is that, since the ARG is a succinct representation of the observed genetic variation, testing clades in the ARG rather than individual SNPs is computationally efficient (and, in the case of Tests 1A and 2, can be performed only once for a genetic dataset and yield results that are applicable to many phenotypes).

The use of WGS data and dense imputation almost resolves the challenge of phantom epistasis by ensuring that most additive causal variants are either directly genotyped or accurately inferred. This leaves phantom epistasis a concern primarily for variants that are poorly imputed or structurally complex. However, this approach is contingent on the availability of large WGS datasets and imputation panels, which are rarely available for species other than humans. Our method therefore provides a robust (and computationally efficient) alternative for epistasis testing in situations where dense imputation is not feasible, such as studies in non-human organisms or when only a relatively small WGS sample is available. While most ARG methods have been tailored for use with human data, this presents a fruitful avenue for future research as availability of sequencing data expands across the tree of life.

## 5 Data availability

Code implementing the methods is publicly available at github.com/a-ignatieva/spectre. Code used to run simulations and produce the figures, and the resulting files, are publicly available at github.com/a-ignatieva/spectre-paper. We used the 1KGP ARG reconstructed using Relate from Speidel et al. (2019). Summary statistics for Hemani et al. (2014) SNPs were obtained from Table 1 of the replication analysis in Wood et al. (2014).

## 6 Acknowledgements

LAFF was supported by the Wellcome Trust [22334/Z/21/Z]. For the purpose of Open Access, the authors have applied a CC BY public copyright licence to any Author Accepted Manuscript version arising from this submission.

## Supplementary Information

### S1 Supplementary methods

#### S1.1 Summary of notation

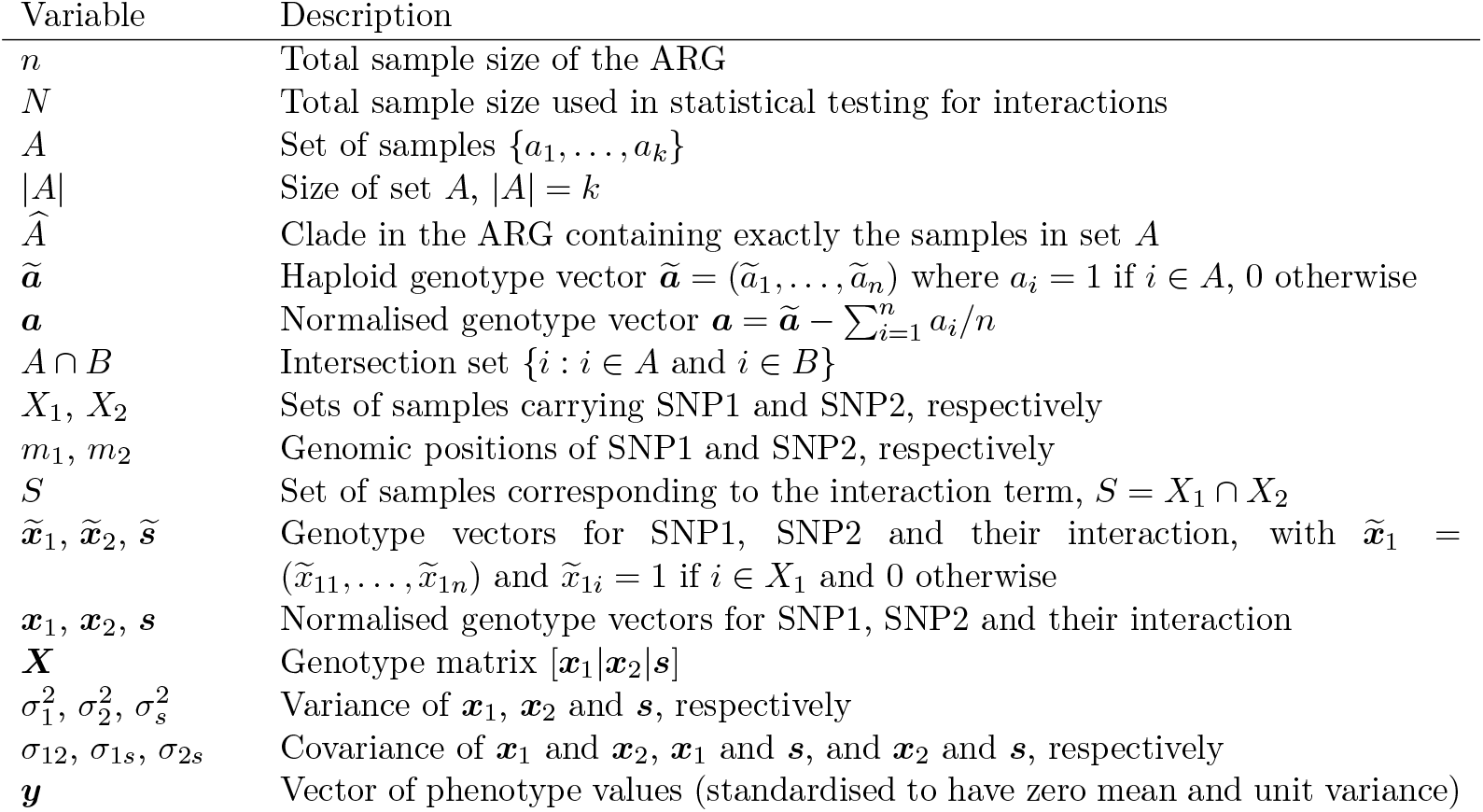

Note that, for readibility, in the main text we do not distinguish between unnormalised and normalised genotype vectors (since this does not cause ambiguity in the narrative). Without loss of generality, all variables (including *s*_*i*_) are assumed to have mean zero. This simplifies derivations as it allows us to omit and ignore intercept terms.

#### S1.2 Probability that a set of samples forms a clade in a local tree

Consider the probability that the target set *S* forms a clade in a random coalescent tree with *n* samples. This requires that the samples in *S* coalesce together before they coalesce with any of the other samples, giving the recursions

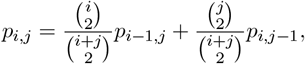

where *i* is the number of lineages currently in the tree subtending *S, j* is the number of lineages currently in the tree *not* subtending *S*, and *p*_*i,j*_ is the probability that *S* forms a clade given *i* and *j*.

The boundary conditions are *p*_1,*j*_ = 1 ∀*j* and we solve for *p*_|*S*|,*n*−|*S*|_. This can be solved explicitly:

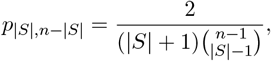

see, for instance, Hein et al. (2004) [p. 84, eq. (3.26)]. For large *n*, unless |*S*| (or *n* − |*S*|) is small, this probability is very small.

However, we know that *S* is a subset of *X*_1_, which forms a clade 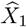 at position *m*_1_. So the relevant probability of interest is instead 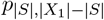, which is larger than *p*_|*S*|,*n*−|*S*|_. In fact, in some *region* around *m*_1_, this probability will be elevated (before the local trees are broken up by recombination). The same argument holds for the other target sets. Notice that this does not depend on *m*_1_ and *m*_2_ being close together. However, if *m*_1_ and *m*_2_ *are* close together, the probability that *S* forms a clade will be much higher, since *S* a subset of *X*_1_ and a subset of *X*_2_,which both form clades within the region [*m*_1_, *m*_2_].

#### S1.3 Statistical conditions for phantom epistasis

In practice, it is enough for a set *Z* which is sufficiently highly correlated with *S* to form a clade 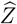 in order for phantom epistasis to emerge. We now derive the exact conditions that such a set must satisfy.

##### S1.3.1 The true model

Suppose that we have data on a sample of *N* individuals, indexed by *i*, for whom we observe a quantitative phenotype *y*_*i*_. We assume the same theoretical setup considered by de los Campos et al. (2019) and so examine the effects of variation at three loci, denoted by *z*_*i*_, *x*_1*i*_ and *x*_2*i*_, on this phenotype. For clarity, we first re-derive the results of de los Campos et al. on the least squares estimator of the interaction coefficient before considering the conditions under which we expect phantom epistasis to occur.

The first variant, *z*_*i*_, is assumed to be the only ‘causal locus’: together with a zero-mean error term *δ*_*i*_ with constant variance for all individuals, it completely determines the phenotype:

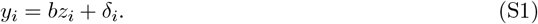

From this theoretical setup, it is clear that there is no epistasis affecting *y*_*i*_. However, we assume that the true causal locus *z*_*i*_ is not observed; instead, we only observe *x*_1*i*_ and *x*_2*i*_, two variants that may be correlated with *z*_*i*_.

##### S1.3.2 The model we fit

We fit the following linear regression model which allows for an epistatic relationship between *x*_1*i*_ and *x*_2*i*_ with an effect on the phenotype:

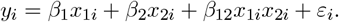

Since the true additive effect *z*_*i*_ is unobserved, fitting the model above may suggest an epistatic relationship between *x*_1*i*_ and *x*_2*i*_ if the estimate for the coefficient *β*_12_ is significant.

We will now derive the probability of obtaining a significant estimate for *β*_12_, which will depend on the partial covariance of the simultaneous occurrence of variants *x*_1*i*_ and *x*_2*i*_ (after conditioning on *x*_1*i*_ and *x*_2*i*_) and the true additive effect *z*_*i*_.

##### S1.3.3 Notation

To simplify notation, we define auxiliary variables *s*_*i*_ := *x*_1*i*_*x*_2*i*_ and *β*_*s*_ := *β*_12_, and so write the model to be estimated as follows:

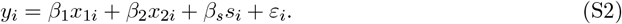

To aid with our derivations below, we also write the model in an equivalent matrix formulation:

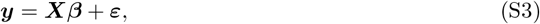

where ***y*** = (*y*_1_, …, *y*_*n*_) (and similarly for ***ε***), the genotype matrix is

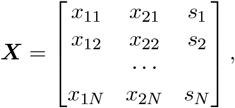

and ***β*** = (*β*_1_, *β*_2_, *β*_*s*_).

##### S1.3.4 Assumptions

We make four additional assumptions. First, all variables are independent and identically distributed across individuals. Second, the variances of, and covariances between, all the independent variables *x*_1*i*_, *x*_2*i*_, *s*_*i*_ and *z*_*i*_ exist and are finite. Third, there is no multicollinearity between the variables *x*_1*i*_, *x*_2*i*_ and *s*_*i*_; in other words, the matrix ***X*** is full rank. This implies, for instance, that SNP1 and SNP2 are not nested in the genealogy (since then we would have *x*_2*i*_ = *s*_*i*_ if SNP2 is on a branch subtended by the branch on which SNP1 is located, for example). Finally, we assume zero correlation between the true error term *δ*_*i*_ and all independent variables in equation (S2) as well as the true causal variant *z*_*i*_, i.e., 𝔼 (*δ*_*i*_*z*_*i*_) = 𝔼 (*δ*_*i*_*x*_1*i*_) = 𝔼 (*δ*_*i*_*x*_2*i*_) = 𝔼 (*δ*_*i*_*s*_*i*_) = 0 for all *i*.

The second and third assumptions are realistic in genetic applications: Bernoulli or binomial random variables always have finite variances and only non-identical genetic variants that segregate in the sample are considered, guaranteeing no multicollinearity. Note also that, while the model above assumes that the phenotype is influenced by only one genetic locus, we could derive the same results while including an additional polygenic component (e.g., 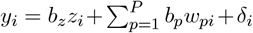) as long as all these additional genetic variants (which would then be part of the error term in the model as it is stated) are uncorrelated with *z*_*i*_, *x*_1*i*_ and *x*_2*i*_ so that the fourth assumption holds. The model is therefore more general than it may appear at first, with the key assumption being that there is only one causal genetic locus in the haplotypic block under consideration.

##### S1.3.5 Asymptotic behaviour of the least squares estimator of *β*

We first emphasise that the true causal locus *z*_*i*_ is not part of the set of independent variables included in our regression (as it is not observed), which implies that it is effectively part of the error term. In other words, we may rewrite the model in equation (S2) as follows:

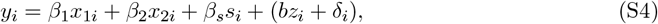

where by assumption we have *β*_1_ = *β*_2_ = *β*_*s*_ = 0 and 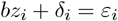

From standard results for least squares estimators of linear models (see, e.g., Wooldridge (2010) [§4.2.1]) we know that the presence of a variable in the error term that has a non-zero effect on the dependent variable renders the estimator of the effect of an included regressor inconsistent as long as this variable is correlated with that regressor. It is this fact – that the least squares estimator may not converge to the true value for the parameter of interest *β*_*s*_, which is zero, even as the sample size becomes infinitely large – that can lead to a false-positive interaction being detected if *z*_*i*_ is correlated with *s*_*i*_ controlling for *x*_1*i*_ and *x*_2*i*_. Note that we cannot check empirically whether such correlations are non-zero as we do not observe *z*_*i*_.

The least squares estimator of the coefficients in the original model as written in (S3), denoted 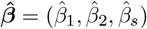 is computed as follows:

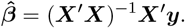

The true model given in equation (S1) is in matrix form ***y*** = *b****z*** + ***δ***. To aid in our derivations below, we add ***Xβ*** (which is zero by assumption) and so have ***y*** = ***Xβ*** + *b****z*** + ***δ***. Plugging in this expression for the true value of ***y*** into the formula for the estimator, we get:

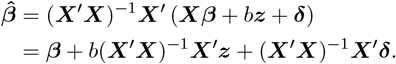

We wish to derive the asymptotic behaviour of 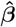. Slutsky’s theorem allows us to write its limiting expression as follows (where 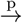 denotes convergence in probability and plim its limit):

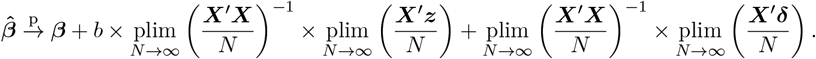

We can now apply the weak law of large numbers, which ensures that sample means converge to expectations as *N* →∞. Defining a matrix ***T*** as follows:

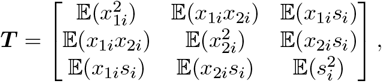

we have:

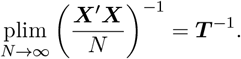

Similarly,

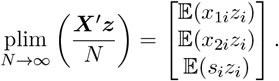

And since we assume no correlation between *δ*_*i*_ and *x*_1*i*_, *x*_2*i*_, *s*_*i*_, we have:

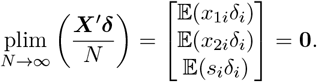

Plugging in these limits, we obtain:

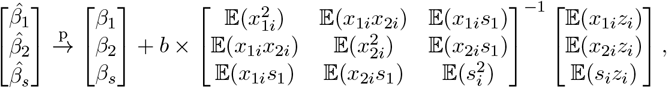

which is equivalent to equation (3) in de los Campos et al. (2019).

Note that the term multiplying *b* is by definition the coefficient vector in the following population linear model, or linear projection,^1^ of the unobserved variant *z*_*i*_ on the observed variants *x*_1*i*_, *x*_2*i*_, *s*_*i*_, which we write as:

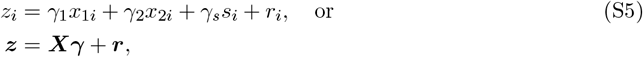

where *r*_*i*_ are the residuals. The vector of population coefficients in this model is:

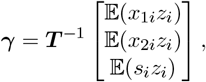

exactly as above.

Focusing now on the estimator of the coefficient of the interaction term 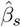 and noticing the connection with the linear projection, we obtain a well-known result: the limit of the least squares estimator is the true value *β*_*s*_ with the added bias of the true effect of the omitted variant *z*_*i*_ which is weighted by the coefficient of *s*_*i*_ in a multiple regression model of *z*_*i*_ on *s*_*i*_ and all other regressors, in this case *x*_1*i*_ and *x*_2*i*_:

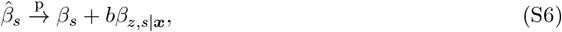

where *β*_*z,s*|***x***_ denotes this multiple regression coefficient. This may also be written as:

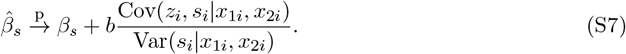

In this example, the interaction *s*_*i*_ has no effect on the phenotype (i.e., *β*_*s*_ = 0) and therefore the estimate of its effect will be driven solely by the tagged effect of the omitted variable. This estimate will tend to be larger the larger the effect (in absolute size) of the true locus and (loosely) the stronger its correlation with the interaction term of interest after accounting for the main effects.

Writing the linear projection of *z*_*i*_ on *x*_1*i*_, *x*_2*i*_, *s*_*i*_ also enables us to derive two more useful facts about the least squares estimator in this context. Plugging in the formula for *z*_*i*_ in equation (S5) into the regression model in equation (S4), we obtain:

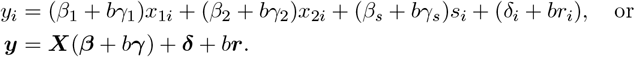

This rewritten model satisfies standard assumptions for least squares estimators: ***X*** is full rank and the error term (comprised of *δ*_*i*_ and *br*_*i*_, where *b* is a constant) is uncorrelated with all regressors and has constant variance. Therefore we can directly invoke known results (see, e.g., Wooldridge (2010) [p. 59, Theorem 4.2]) to establish the asymptotic Normality of the estimator:

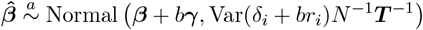

and the consistency of the usual estimator of the variance of 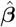 :

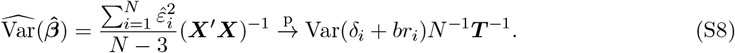

##### S1.3.6 Deriving the probability of false positive interactions

Having derived the asymptotic behaviour of both the estimator of the coefficient 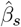 and its standard error, we now consider the asymptotic behaviour of the test statistic that is used to assess a possible effect of the interaction term *s*_*i*_ on the phenotype.

A standard *t* -test of the null hypothesis *β*_*s*_ = 0 is based on the following test statistic:

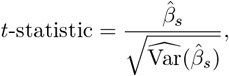

which for large samples can be compared to a chosen quantile of the standard Normal distribution.

For a chosen level of significance *α*, and defining 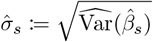 for conciseness, we reject the null hypothesis in a two-sided test if and only if:

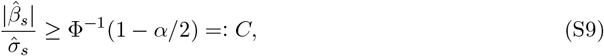

where Φ(·) is the cumulative distribution function of the standard Normal.

The question we seek to answer is: for an interaction term *s*_*i*_ with no effect on the phenotype but which tags an unobserved third variant *z*_*i*_ with a non-zero effect *b*, under what conditions (on the effect size of *z*_*i*_ and on the relationship between *s*_*i*_ and *z*_*i*_) do we expect to *incorrectly* reject the null hypothesis of no effect for *s*_*i*_? This hinges on whether the inequality in equation (S9) is satisfied.

We have seen that 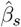 is asymptotically Normally distributed with mean *bβ*_*z,s*|***x***_ and a variance which is estimated consistently through the usual approach, which implies that the test statistic will be asymptotically Normal with mean 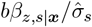 and unit variance. The probability *P* of incorrectly rejecting the null hypothesis is then equal to the probability that a random draw from such a Normal distribution exceeds the significance threshold *C*:

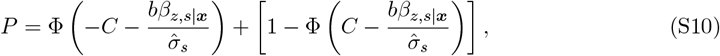

where the first term on the right hand side corresponds to the shaded area on the left in Figure S1a and the second term to the area on the right in the Figure.

Equation (S10) has four variables (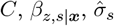 and *b*) which together determine the probability of interest *P*. Conversely, fixing the probability *P* and three of these variables yields a value for the fourth one. In practice, *C* is determined by the significance level of the original interaction test whose results we assess and is therefore given. Moreover, *β*_*z,s*|***x***_ can be readily estimated for each variant *z* under consideration from the external dataset on which the ARG is built.

Turning now to the third variable 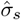, since this is the standard error of the interaction coefficient in the regression model it could also be taken from the summary statistics of the interaction test. However, we propose instead to approximate it in a manner that is both highly accurate and which makes this approach independent of the particular phenotype under analysis (the advantages of this will be discussed below). From equation (S8), we see that 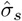 is computed as follows:

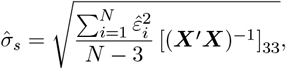

where (***X***^′^***X***)^−1^ denotes the bottom right (third row, third column) element of the matrix

(***X***^′^***X***)^−1^, which corresponds to the regressor *s*_*i*_. Since the phenotypes on which GWASs are run are generally highly polygenic, with hundreds or even thousands of independent associations each with a very small effect, the three regressors in the model given in equation (S2) will jointly account for only a very small percentage of the variance of the phenotype. This in turn implies that the variance of the residuals 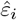 will be approximately equal to the sample variance of the phenotype itself. Therefore we can very accurately approximate 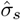 as follows:

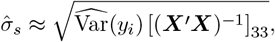

where 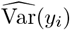 is the sample variance of the phenotype. In the derivations above, we could have assumed without loss of generality that the phenotype had been standardised and so had unit variance, in which case we would have 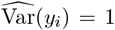. However, we need n ot make this additional assumption. If we simply approximate 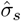 as 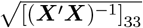, the term 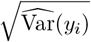 will implicitly divide *b* in equation (S10). Therefore the values of *b* that we will obtain through this equation, as we explain in the next paragraph, will be rescaled so that their units are standard deviations of the phenotype and are thus readily interpretable independently of the scale on which the original phenotype is measured – this is a first advantage of employing this approximation. Note that, like *β*_*z,s*|***x***_, [(***X***^′^***X***)^−1^]_33_ can also be easily computed from the external dataset from which the ARG was constructed.

We have seen how the values for three of the four variables in equation (S10) can be set and now turn to consider the fourth variable *b*. This variable is fundamentally unknowable as we cannot by definition estimate the effect of an unobserved variant. We therefore propose to set a range of probabilities *P* of incorrectly rejecting the null and, for each of these probabilities, compute the value of *b* that exactly yields that probability given *C, β*_*z,s*|***x***_ and 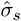 (with larger absolute values of *b* increasing this probability). One can then assess whether the smallest value that the effect of a potential third variant needs to take to lead to an inferential error with a certain probability is small enough that the existence of such a variant is plausible and so warrants concern.

**Figure S1:**
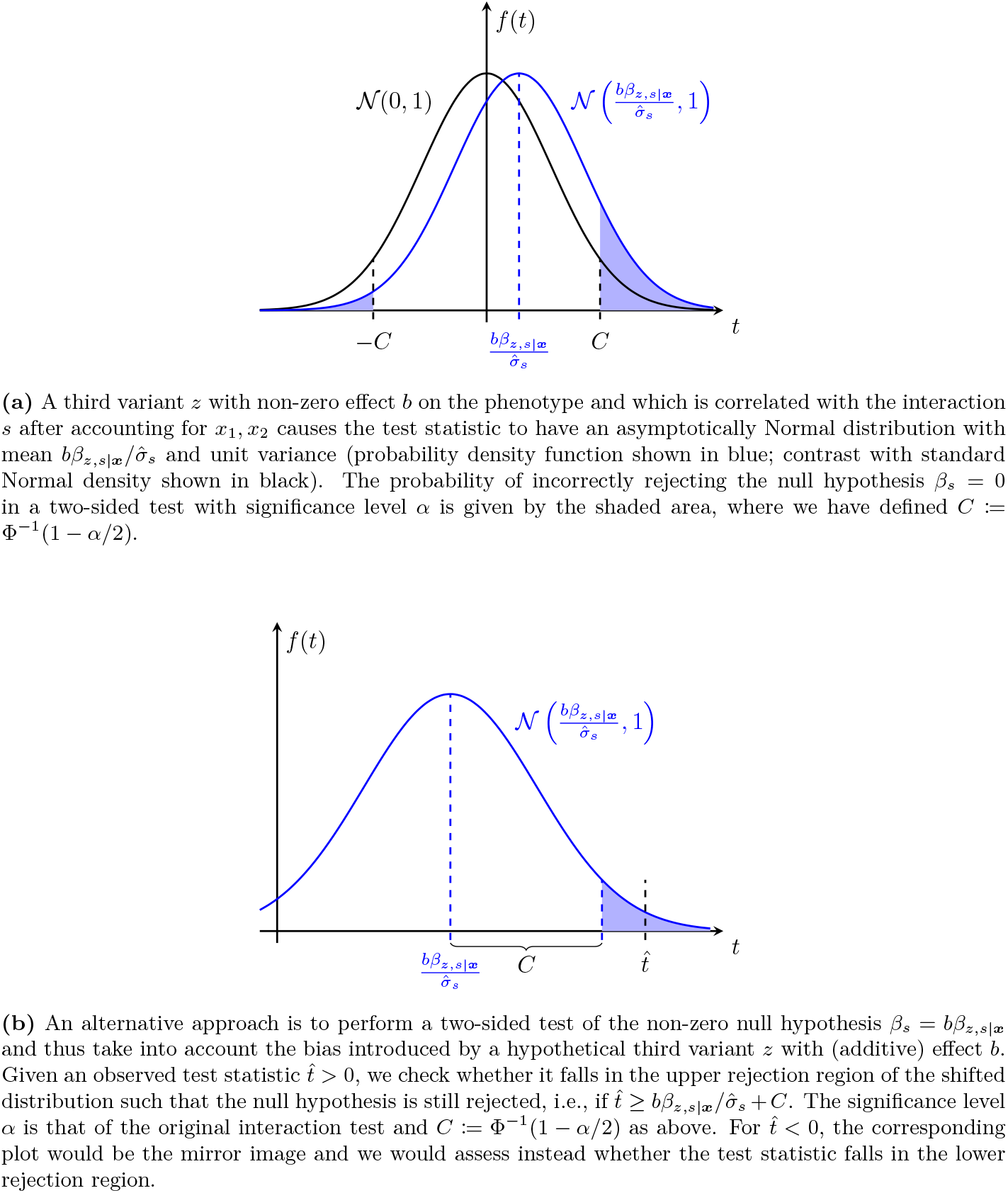
Two alternative approaches to assessing the reliability of interaction signals given the possible presence of a third variant with non-zero effect on the phenotype.

An alternative approach is to assume that there is a problematic third variant, adjust the null hypothesis to account for the expected bias in the inferred coefficient 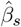 (which we derived above) and then test this adjusted hypothesis that *β*_*s*_ = *bβ*_*z,s*|***x***_. Intuitively, if 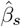 is very large (in absolute value) then we would still reject the adjusted null if the additive effect of the third variant *z* and its partial correlation with *s* are not too strong. We then ask, how large can the bias of the test statistic 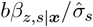 be such that we still reject this non-zero null hypothesis?

We consider a two-sided test of the null hypothesis *β*_*s*_ = *bβ*_*z,s*|***x***_ against the alternative *β*_*s*_ ≠*bβ*_*z,s*|***x***_ with the same significance level *α* as the original interaction test. For a positive observed value of the test statistic, which we denote by 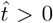, we check whether this statistic falls in the upper rejection region of such a test, i.e., if 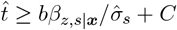 where *C* := Φ^−1^(1 − *α/*2) as above (Figure S1b). (If instead 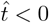, we would assess whether the test statistic falls in the lower rejection region, i.e., if 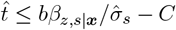.)

In practice, given the significance level of the original test, the estimated coefficient *β*_*z,s*|***x***_and the standard error 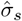 (set to 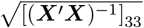 as before), we compute the value *b* that makes the boundary of the rejection region exactly equal to the observed test statistic so that the null hypothesis is just rejected. For 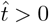 and *β*_*z,s*|***x***_ *>* 0, as well as for 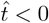 and *β*_*z,s*|***x***_ *<* 0, this gives a positive upper bound for *b*, with smaller values moving the rejection region boundary away towards zero. Conversely, for 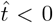 and *β*_*z,s*|***x***_ *>* 0, or for 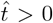 and *β*_*z,s*|***x***_ *<* 0, we would obtain instead a negative lower bound for *b*. We can then interpret the results as meaning that the interaction identified would still be significant in a two-sided hypothesis test with the same significance level as the original test as long as the effect *b* of the correlated clade satisfies this bound. (By focusing on the rejection region farthest from zero, we only reject the null for observed values of 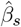 which are larger in absolute value than the expected bias *bβ*_*z,s*|***x***_ for this coefficient. If we considered also the lower rejection region, our range of ‘allowed’ values for *b* would also include values so large in absolute value that the null hypothesis would be rejected because the observed test statistic was too close to zero, which would be unnatural.)

These two approaches are complementary but distinct as they use different sets of information. The first approach does not use the estimated coefficient 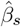 or the resulting test statistic, and is thus independent of the phenotype under analysis (we ensured this was the case by approximating 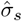 without using phenotypic information – this general applicability independent of phenotype is a second advantage of the proposed approximation). It simply asks whether two variants can reliably be tested for the effect of their interaction (on any phenotype, provided that it is highly polygenic – or, more precisely, that the variants in question account for very little phenotypic variance – such that the approximation to 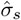 is accurate) with some confidence that a possible interaction signal would not be caused by a hidden third variant, and indeed can be run prior to interaction testing. Given a genetic dataset for which multiple phenotypes are available, this approach could be applied to all pairs of variants within a certain distance from each other to identify pairs that can be tested for interaction; then in a second step we could test whether each of these pairs of SNPs had a significant interaction for each phenotype separately.

In contrast, the second approach does use the estimated test statistic and so requires interaction testing to be performed first and is phenotype specific. It can be thought of as qualifying the interaction signals by asking how robust these are to the presence of hidden third variants: everything else being equal, stronger signals (with a smaller *p*-value) will be able to withstand third variants with a larger additive effect; however, this robustness will decrease for variants with a highly-correlated neighbouring clade.

#### S1.4 Searching for clades correlated with a target set

For each clade 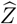 of the ARG within ±1cM of each SNP in the interaction pair under consideration, we can calculate the corresponding normalised genotype vector ***z*** and regress this vector on ***x***_1_, ***x***_2_ and ***s*** to obtain 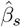 (and repeat this replacing ***s*** with each of the other target set genotype vectors). This is the basis for the procedure described in Methods, Section 2.4.1: the relevant value of *b* that can be problematic as just described is then based on the largest value of 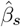 across all clades and target sets.

**Figure S2:**
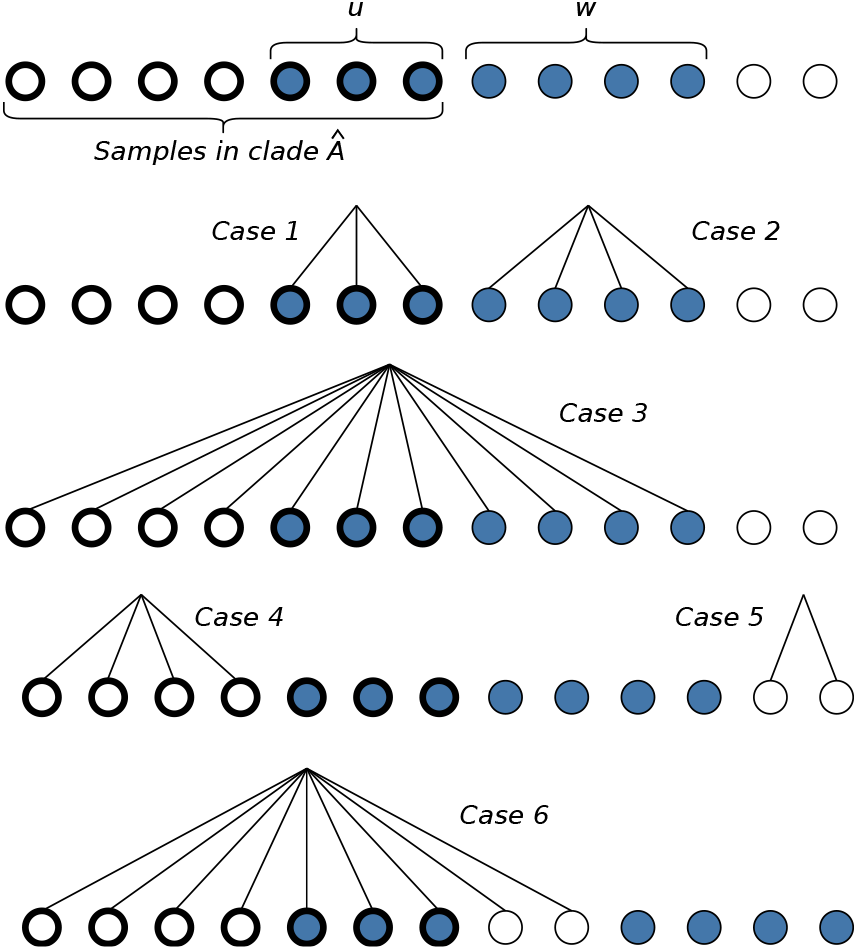
Samples of an ARG (ARG itself not shown) of size *n* = 14. Samples in clade *Â* shown with bold outline; samples in set *S* shown in blue. Lines connect samples that could possibly form a clade 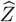 in each considered case. Note that since we condition on *A* itself forming a clade, we cannot observe a clade formed of some samples from within *Â* and some samples from outside *Â* as this would violate that assumption (for instance, in the scenario shown, the set *S* itself cannot form a clade — only a set correlated with *S* can).

#### S1.5 Evidence against existence of clades correlated with a target set

Let *S* be the set of samples in the ARG corresponding to the interaction genotype vector ***s*** (that is, the set of samples that carry both SNP1 and SNP2). Suppose clade *Â* appears in some local tree along the genome, with |*S* ∩ *A*| = *u*, and let *w* = |*S*| − *u* (as illustrated in Figure S2, top line). We would like to check whether *Â* provides evidence against the existence of another clade 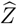 (within the genomic span of *Â*) such that the corresponding normalised genotype vector ***z*** would satisfy (S9).

Since, in realistic scenarios, ARG reconstruction is not perfect and there is potentially a large amount of uncertainty about the topology of each local tree, we condition only on the fact that the samples in set *A* form a clade *Â*, that is, we use only the most relevant information provided by the ARG on clade existence and genomic span (rather than conditioning on all of the information contained in the graph, such as mutation ages). Then there are six possible scenarios to consider which maximise the correlation between a possible clade 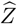 and the set *S*, shown in Figure S2. Cases 1–3 (resp. 4–6) correspond to maximising the positive (resp. negative) correlation between *Z* and *S*, when 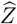 is a subclade (Cases 1 and 4), superclade (Cases 3 and 6) or disjoint from *Â* (Cases 2 and 5).

Taking each case in turn, for a fixed probability *P*, we calculate *b* from (S10), and take the minimum (and we repeat this procedure for each target set). Then, at each position of the genome, the maximum such value over all tested clades that span that position is informative of whether phantom epistasis can be effectively ruled out at that position. That is, we check whether the topology of the ARG implies that the samples in each target set are too well-separated for any problematic clade (one that is highly correlated with a target set) to be feasible. This is the basis for the procedure described in Methods, Section 2.4.2.

Note that, to limit the number of cases considered for each clade to six, we make a key simplifying assumption that *Z* being highly correlated with *S* is more likely to cause (S9) to be satisfied, which we expect to hold in practice unless *S* is small. Our simulation studies support this assumption: in cases of phantom epistasis, the mean correlation between the most significant explanatory clade and a target set was 0.32.

---

**Figure S3:**
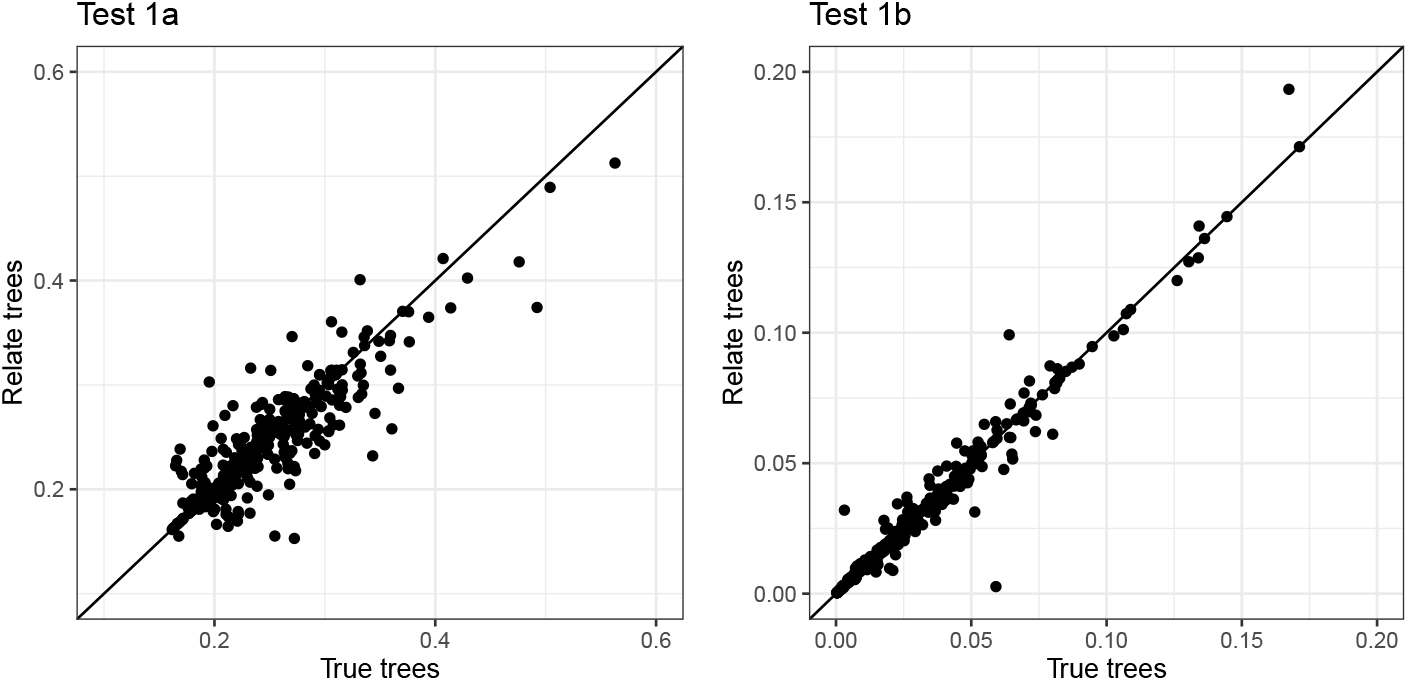
Values of *b* from Test 1A (left panel) and 1b (right panel) for true trees (*x*-axis) and Relate trees (*y*-axis), demonstrating close agreement.

## Footnotes

1 A unique linear projection of this form is guaranteed to exist since the matrix ***X*** is non-singular; see, e.g., Wooldridge (2010) [§2.3].

